# A maximum likelihood estimator for quartets under the Cavender-Farris-Neyman model

**DOI:** 10.1101/2024.08.16.608292

**Authors:** Max Hill, Jose Israel Rodriguez

**Affiliations:** Department of Mathematics, University of California, Riverside, Riverside, USA, 92521; Department of Mathematics, Department of Electrical and Computer Engineering, University of Wisconsin — Madison, Madison, USA, 53703

## Abstract

We present the Julia package `FourLeafMLE.jl`, which solves a global optimization problem to perform statistical inference using DNA sequence alignment data for quartets under the Cavender-Farris-Neyman phylogenetic model.

## 1 Introduction: tree topologies and the tree of life

One of the core motivations in evolutionary biology is to reconstruct the *tree of life* that describes how species have evolved over time. A common approach is to use DNA sequence alignment data from present-day taxa to infer a *phylogenetic tree*, understood as a leaf-labelled tree with edges representing ancestral populations, vertices representing common ancestors, and leaves representing extant taxa. The edge weights, or *branch lengths* represent some measure of evolutionary time (usually expected number of mutations per site), and the tree *topology* describes clade structure of the taxa. For instance, Figure 1a has four leaves that are labeled in such a way to indicate species 1 and 2 have a more recent common ancestor with one another than with species 3 or 4.

**Figure 1.**
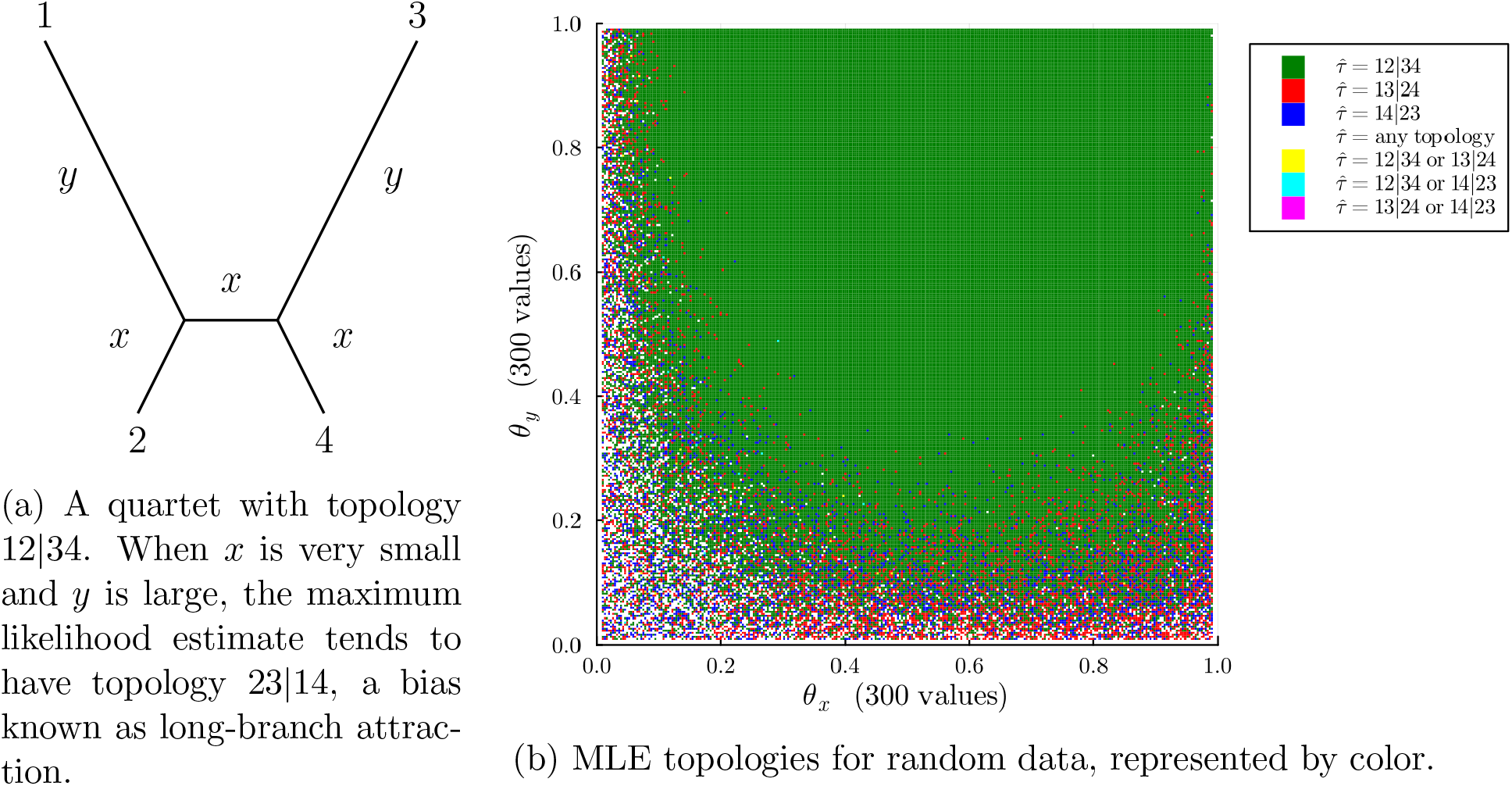
A visualization of 90, 000 inferred tree topologies (right) for random data consisting of 1000 bp drawn according to the CFN model on trees of the form given on the left. The reddish bottom right corner of (b) provides a new visualization of the long-branch attraction phenomenon.

A phylogenetic tree thus represents a hypothesis about the evolutionary history of a set of taxa; the statistical inference problem is to infer the tree topology and branch lengths from a DNA sequence of fixed length. Our implementation solves polynomial systems to perform this inference.

## 2 Maximum likelihood estimation & quartet methods

A standard approach to inferring a phylogenetic tree is to use maximum likelihood estimation. By summarizing data as a vector (*u*_1_, …, *u*_*n*_) of counts, the goal is to maximum the likelihood of the data. In other words, we seek the value of *θ* that maximizes 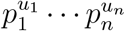: where *p*_*i*_ is the probability of observing event *i* and each *p*_*i*_ depends on the parameters *θ* of the model. The maximum likelihood problem addressed in `FourLeafMLE.jl` involves estimating the branch lengths and unrooted tree topology of a 4-leaf tree from sequence data of a fixed length *k* generated according to the Cavendar-Farris-Neyman (CFN) model [12].

Considerable research has focused on understanding the properties of maximum likelihood estimation. Even in the simplest cases of 3- and 4-leaf trees, the problem exhibits substantial complexity, with a general solution known for 3-leaf trees, but not 4-leaf trees [1, 4, 6, 7, 8, 9, 11, 16]. The 4-leaf case considered here is of special interest first due to the popularity of quartet-based inference methods for inferring both phylogenetic trees and networks [15], and second because it is the simplest case in which the phenomenon of *long-branch attraction* can be observed, a form of estimation bias which is only partially understood [2, 14].

Two challenges in optimizing the likelihood function are its non-convex nature and the presence of numerous boundary cases during optimization. Moreover, there can be multiple distinct maximizers of the likelihood function, and the maximizers can have branch lengths which are infinitely long or zero [5, 13]. This package resolves these two obstacles for the CFN model by using computer algebra and the theory of maximum likelihood (ML) degrees [10]. One feature distinguishing this software is that it implements a framework to characterize cases when the tree estimates have infinite or zero-length branches, which are cases where the MLE may not take the form of a connected, 4-leaf binary tree. We believe this addition may be useful in understanding the ways that maximum likelihood inference can fail, and as a first step leads to Figure 1.

## 3 Running the maximum likelihood estimator

Our main contribution puts algebra into practice by implementing a custom tailored solver for algebraic statisticians. Given data in the form of a site frequency vector consisting of 8 entries giving the frequencies of the patterns (*xxxx, xxxy, xxyx, xxyy, xyxx, xyxy, xyyx, xyyy*), the main functionality is to return a list of global maximizers of the likelihood function. The last entry of the output includes a description of the tree corresponding to the maximizer. There are an additional 9 types of cases where the output does not correspond to a binary 4-leaf tree; more details on interpreting the output for such cases is included in the documentation.

`SITE_PATTERN_DATA = [212, 107, 98, 115, 114, 89, 102, 163] fourLeafMLE(SITE_PATTERN_DATA)`

`## Output`

`1-element Vector{Vector{Any}}:`

[-2036.1212979788797, “R1”, [1],

`[0.1380874841, 0.46347429951, 0.5231552324, 0.3975875363, 0.6835395124],`

`“θ1, θ2, θ3, θ4, θ5”,`

`“binary quartet with topology τ=12|34 and edge parameters (θ1,…,θ5) = (0.1380874841,…,0.6835395124)”]`

For complete details on the code, see our Julia package^1^, but now we give a broad overview about our software. The CFN model is a parameterized semialgebraic set in R^8^ with dimension 5. We optimize the likelihood function by partitioning the semialgebraic set according to boundaries from the parameterization, similar to the approach in [1] for a different model. There are 10 different boundaries up to symmetry. All but one of them have ML degree less than or equal to two. The last one has ML degree 92 while the main component of the semialgebraic set has ML degree fourteen.^2^ We optimize the likelihood function on each of the boundary cases by solving the likelihood equations [10] by using analytic expressions when the ML degree is less than or equal to two and `HomotopyContinuation.jl` [3] to compute the critical points otherwise.

## 4 Put into practice: Visualizing the geometry of data

Our method is fast. On a desktop machine with 12 i5-10400 CPUs (2.90GHz), the command `FourLeafMLE.jl` solves on average 11 global optimization problems per second. This is sufficiently fast to create high-quality visualizations of the geometry of the maximum likelihood problem — instructions are printed by the command @doc make classification plot. Figure 1b presents a grid of 90, 000 points, each representing a choice of Hadamard edge length parameters *θ*_*x*_, *θ*_*y*_ (see [12]) for the tree shown in Figure 1a. For each point, maximum likelihood estimation was performed using random data (1000 bp) drawn from the model, and the point colored according to which binary quartet topologies were found to be compatible with the maximum likelihood estimates.

## Acknowledgements

We are thankful for the comments by Marina Garrote-López and the anonymous referees that led to improvements of earlier versions of this article. This research was partially supported by the Alfred P. Sloan Foundation. Support was also provided by the Graduate School and the Office of the Vice Chancellor for Research at the University of Wisconsin-Madison with funding from the Wisconsin Alumni Research Foundation.

## Footnotes

1 Available at https://github.com/max-hill/FourLeafMLE.jl.

2 The numbers 14 and 92 were previously reported in [7, Table 2].

## References

[1] E. S. Allman, H. Banos, R. Evans, S. Hosten, K. Kubjas, D. Lemke, J. A. Rhodes, and P. Zwiernik. Maximum likelihood estimation of the latent class model through model boundary decomposition. Journal of Algebraic Statistics, 10(1):51–84, 2019.

[2] J. Bergsten. A review of long-branch attraction. Cladistics, 21(2):163–193, 2005.

[3] P. Breiding and S. Timme. HomotopyContinuation.jl: A Package for Homotopy Continuation in Julia. In International Congress on Mathematical Software, pages 458–465. Springer, 2018.

[4] B. Chor, M. Hendy, and D. Penny. Analytic solutions for three taxon ML trees with variable rates across sites. Discrete Applied Mathematics, 55(6-7):750–758, 2007.

[5] B. Chor, M. D. Hendy, B. R. Holland, and D. Penny. Multiple maxima of likelihood in phylogenetic trees: An analytic approach. Molecular Biology and Evolution, 17(10):1529–1541, 10 2000.

[6] B. Chor and S. Snir. Molecular clock fork phylogenies: Closed form analytic maximum likelihood solutions. Systematic Biology, 53(6):963–967, 2004.

[7] L. D. Garcia Puente, M. Garrote-Lopez, and E. Shehu. Computing algebraic degrees of phylogenetic varieties. Journal of Algebraic Statistics, 14(2):215–231, 2023.

[8] M. Hill, S. Roch, and J. I. Rodriguez. Maximum Likelihood Estimation for Unrooted 3-Leaf Trees: An Analytic Solution for the CFN Model. Bull. Math. Biol., 86(9):Paper No. 106, 2024.

[9] A. Hobolth and C. Wiuf. Maximum likelihood estimation and natural pairwise estimating equations are identical for three sequences and a symmetric 2-state substitution model. Theoretical Population Biology, 2024.

[10] S. Hoşten, A. Khetan, and B. Sturmfels. Solving the likelihood equations. Found. Comput. Math., 5(4):389–407, 2005.

[11] D. Kosta and K. Kubjas. Maximum likelihood estimation of symmetric group-based models via numerical algebraic geometry. Bull. Math. Biol., 81(2):337–360, 2019.

[12] C. Semple and M. Steel. Phylogenetics. Oxford University Press, 2003.

[13] M. Steel. The maximum likelihood point for a phylogenetic tree is not unique. Systematic Biology, 43(4):560–564, 1994.

[14] E. Susko and A. J. Roger. Long branch attraction biases in phylogenetics. Systematic Biology, 70(4):838–843, 02 2021.

[15] T. Warnow. Computational Phylogenetics: An Introduction to Designing Methods for Phylogeny Estimation. Cambridge University Press, 2017.

[16] Z. Yang. Complexity of the simplest phylogenetic estimation problem. Proceedings of the Royal Society of London. Series B: Biological Sciences, 267(1439):109–116, 2000.

